# Effect of temperature and agitation rate variation on the growth of NS0 cell line in monoclonal antibody production

**DOI:** 10.1101/822601

**Authors:** Shamitha Shetty

## Abstract

The quality and yield of the monoclonal antibodies produced in a cGMP environment is heavily influenced by the bioprocess-related parameters which impact the cell growth and metabolism of the mammalian cell cultures. This research report describes a study conducted to examine the effects of varying temperature and RPM set points on viable cell density and viability of NS0 cultures. All cultures were grown in 250 mL shake flasks (working vol. 100 mL). To separately analyze the effects of temperature and agitation rate on NS0 cell metabolism, flask stage cultures were evaluated in triplicates at two cultivation temperatures (36 °C and 38 °C) and two agitation rates (120 RPM and 160 RPM) while controls were maintained at 37 °C and 140 RPM for both the conditions using an incubator. Flasks were sampled every 24 h and analyzed for viable cell density and % viability. Additional data was collected on pH, pO_2_, pCO_2_, osmolality, glucose, lactate, glutamate and glutamine levels in the culture. It was observed that variations in temperature has the greatest effect on viable cell density and viability and varying agitation rates had minimal effect on growth of cultures. A temperature set point of 38 °C is detrimental to the culture growth. The control set points proved to be optimal for this process.

---

Product X is a recombinant, human, IgG_1_λ monoclonal antibody that recognizes human B Lymphocyte Stimulator (BLyS). It inhibits the binding of soluble BLyS to B cells and hence prevents the survival and differentiation of selected B-cell subsets. Product X is used for the treatment of systemic lupus erythematosus (SLE) in adult patients with active, autoantibody-positive SLE who do not respond to standard therapy. Patients with autoimmune diseases like SLE show elevated levels of BLyS cytokine.^1, 2^ This antibody is expressed in the GS-NS0 mouse myeloma cell line using the glutamine synthetase expression system. ^3^ **Figure 1** shows the upstream process that is followed for manufacturing Product X. ^4^ The cell culture process for Product X consists of three stages: S1, S2 and S3. The purpose of the small-scale S1 and S2 seed expansion stages is to produce inoculum of a sufficient viable cell density (VCD) and viability for inoculation of the S3 production bioreactor.

**Figure 1.**
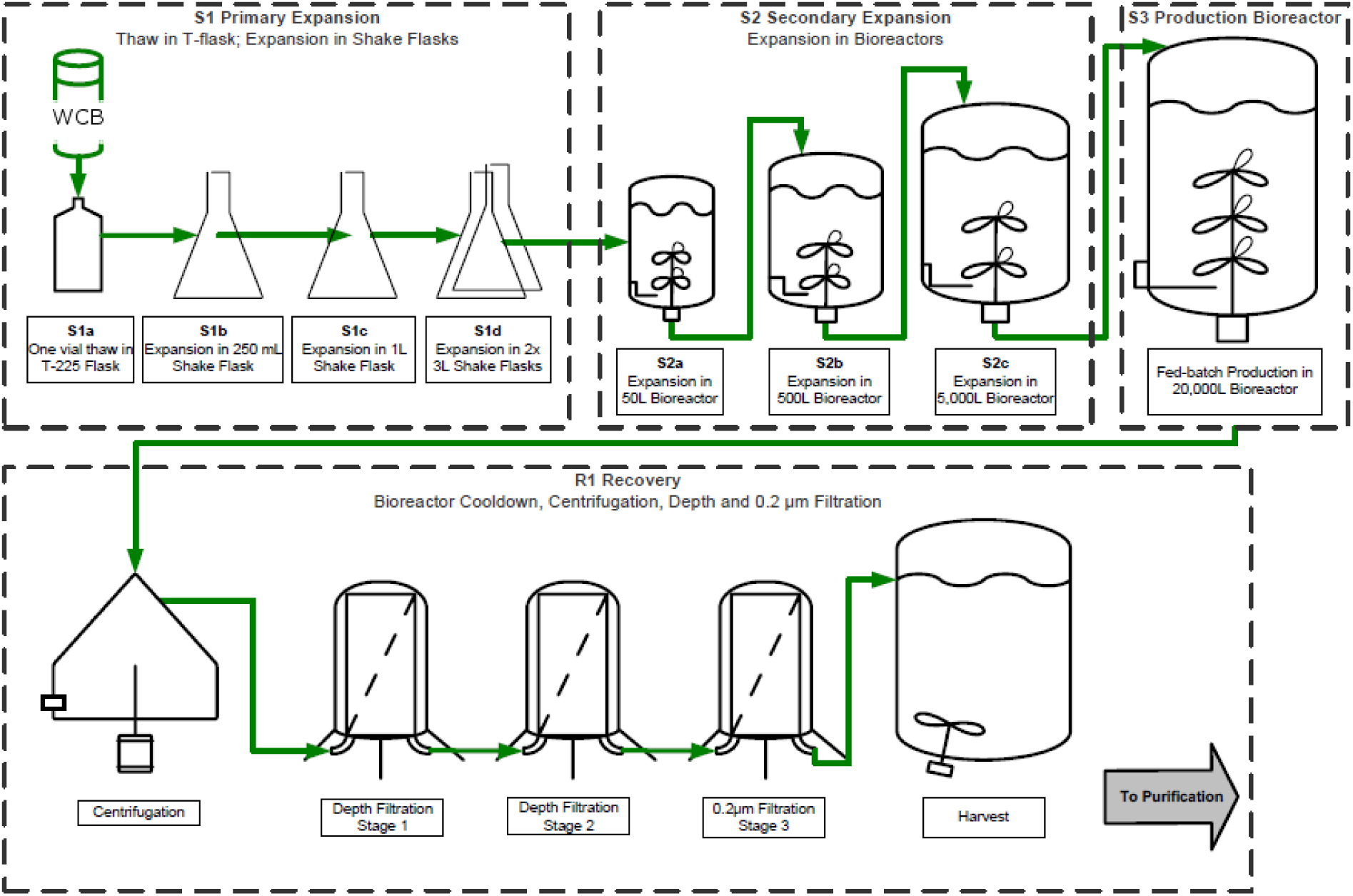
Process Flow Diagram. The Upstream process followed for manufacturing Product X involves stages S1 through Recovery. The Working Cell Bank (WCB) is thawed and inoculated in the 250 mL Shake Flask stage. Further expansion involves the cells being grown in bioreactors until they reach the 20,000 L Production Scale Bioreactor. The cells are then recovered using a series of centrifugation and filtration steps before being harvested and transferred to downstream purification to get the bulk drug substance.

During S1, a vial of the Working Cell Bank (WCB) is thawed into a 125mL shake flask. The culture is expanded in a train of shake flask cultures of increasing scale 125 mL (S1a), 250 mL (S1b), 1 L (S1c), and 3 L (S1d). The S1d culture is used to inoculate the first S2 bioreactor. During S2, cells are further expanded in a train of bioreactors of increasing volume. The S2 train in the large-scale manufacturing facility consists of 50 L (S2a), 500 L (S2b), and 5000 L (S2c) bioreactors. The cell culture continues to grow until it reaches the final cell density range required for transfer to the next step. The purpose of S3 is to produce Product X of sufficient titer and quality for further downstream processing. S3 is a 11 day fed-batch process performed in a 20,000 L bioreactor.

The S3 culture process is completed when harvesting criteria are all simultaneously within their limits (titer, final viability, and maximum VCD) and the material in the production bioreactor is considered Unprocessed Bulk (UPB). The recovery process is initiated at the end of S3 where the culture is clarified by a continuous series of unit operations consisting of stacked-disc centrifugation, depth filtration, and 0.2µM membrane filtration. Following membrane filtration, the supernatant is stored in a chilled holding tank up to 72 h until the purification process begins. Cell free supernatant obtained post recovery process is denoted as Clarified Unprocessed Bulk (CUB). The secreted Product X is recovered from the cell culture production medium and purified using a series of chromatographic and filtration steps. The final drug product is manufactured from the bulk drug substance to produce a single-use, lyophilized or liquid product.

For the scope of the paper, we have focused on the 250 mL Shake Flask (S1b) stage. The incubation conditions for this stage are 37 °C, 5% CO_2_ and 140 RPM agitation rate. It is not known what effect a variation in any of these set points would have on cell growth. The aim of the research is to examine the impact of varying the set points of critical process parameters (CPPs) like temperature and process parameters (PPs) like agitation rate on the growth rate of the cells which is assessed by the VCD and viability of the cells. The growth of cells in control flasks to flasks with high and low set points of temperature and agitation rate will be tested and compared during the experiment.

In recent years, cell culture bioprocessing has seen a tremendous growth in data generation and collection. In modern manufacturing facilities, it is not uncommon to encounter hundreds of process parameters being monitored and acquired automatically every few seconds throughout the entire production train. This enormous volume of data further accumulates across multiple campaigns and at multiple manufacturing sites. Mining these historical data and statistical analysis of the data generated holds promise to gain insights into fluctuations in process performance, uncover hidden characteristics of high-performing cultures, and discern process parameters with pivotal contributions to the overall process performance. ^5, 6, 7^

We have designed an experiment which tests different treatments to the cells for testing the impact of fluctuations in these process parameters on cell culture and collected data throughout the growth stage. We then performed a statistical multivariate and univariate analysis of the data using JMP software to understand the contribution of these parameters to the growth rate over the span of five days.

## RESULTS

### Effect of temperature on cell growth

Based on the data, temperature variations seemed to have a great impact on the VCD and viability of the cells. As shown in **Figure 2**, a least squares model was fitted for the data from all the flasks. The R^2^ value of 1 and 0.92 for VCD (**Fig. 2a**) and %Viability (**Fig. 2b**) respectively suggests that the model is a good fit for the data. It is also seen that all the points lie well along the 45 degree diagonal line which again indicates goodness-of-fit.

**Figure 2.**
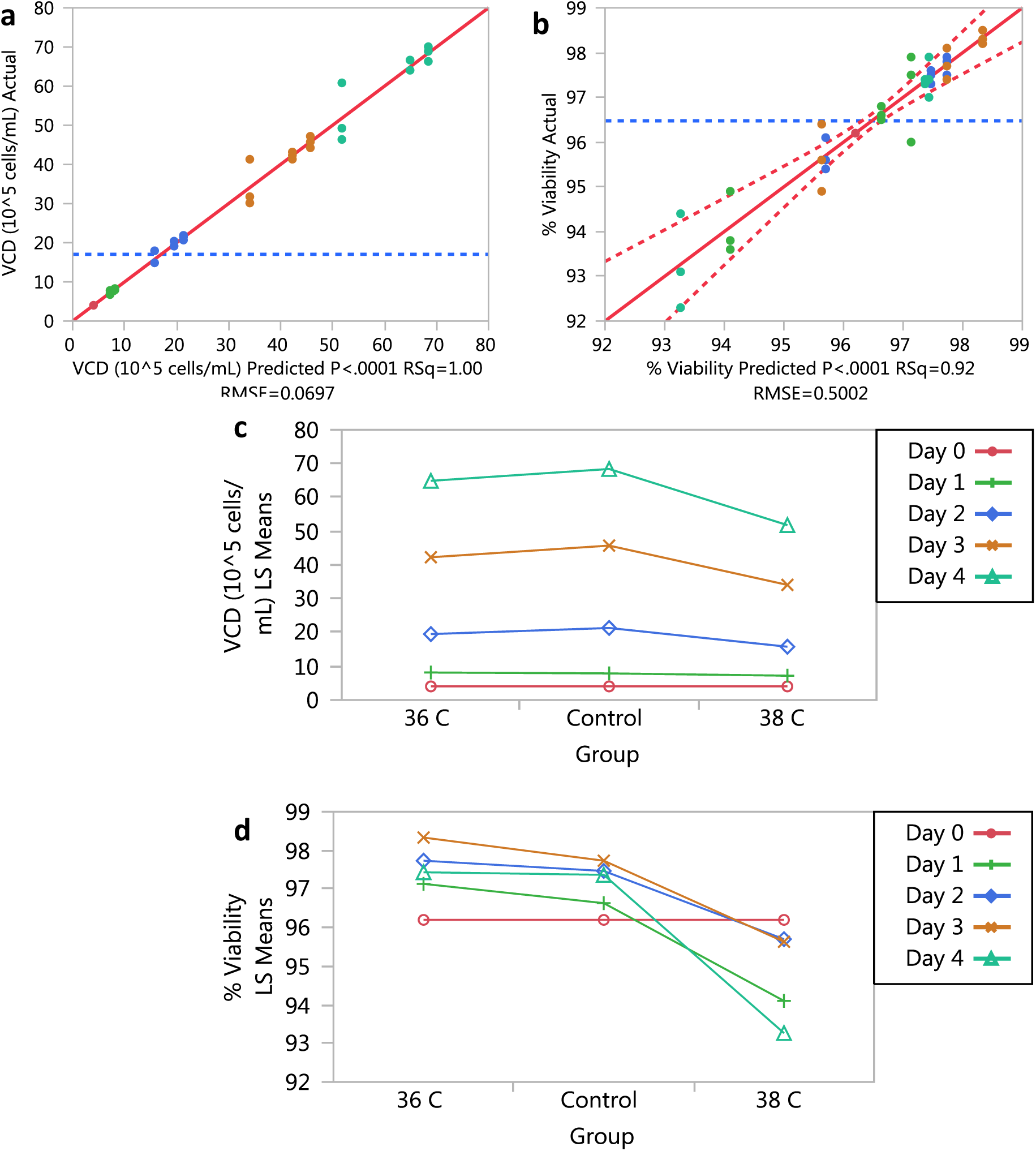
After fitting the linear model using regression analysis in JMP we need to determine if the model fits the data well. The results observed from the analysis can then be used to view any interactions between the responses. **(a,b)** Actual by Predicted plot. The observed values are plotted on the y-axis and the jackknife predicted values of the responses (effects) are on the x-axis for VCD as shown in **a** and %Viability as shown in **b**. This is the leverage plot for the whole model. We can predict the goodness-of-fit for the model by seeing how well all the points lie along the 45 degree line. The closer the value of R^2^ to 1 the better the model fits the data. As both these conditions were satisfied both the models were adjudged to be a good fit for the data. **(c,d)** Least Squares Means (LSMeans) plot. The least squares means for nominal and ordinal main effects and interactions were plotted after fitting the model in JMP. The results showed that in **c**, the VCD levels were most affected by the higher temperature set point. Over the duration of 4 days, the control group seemed to show the highest viable cell density indicating that the control temperature was best suited for the cell growth. Similar results were observed in **d**, but the 36 °C set point flasks seemed to show the highest %Viability as compared to the other two set points indicating that lower temperature was more suited for maintaining the longevity of the culture. At the end of Day 4 however, both the 36 °C and control showed nearly the same values for %Viability. The 38 °C set point proved to be the most unfavorable for cell viability.

Next, the Least Squares values from the model were plotted to view the results of the analysis. It showed the effects of all three temperature set points on the viable cell density across the span of 0-4 Days. It was observed that the flask incubated at 38 °C showed the lowest mean VCD after 4 days and the control flask showed the highest mean VCD (**Fig. 2c**). We observed similar results in the Least Squares Means (LSMeans) plot for %Viability, but the 36 °C set point flasks seemed to show the highest %Viability as compared to the other two set points indicating that lower temperature was more suited for maintaining the longevity of the culture. At the end of Day 4 however, both the 36 °C and control showed nearly the same values for %Viability. The 38 °C set point again proved to be the most unfavorable for cell viability. The flask incubated at 38 °C reached a maximum VCD of 60.84 × 10^5^ cells/mL as compared to a VCD of 66.67 × 10^5^ cells/mL in the 36 °C flask and 70.08 × 10^5^ cells/mL in the control flask. After 4 days, the 38 °C flask had a maximum viability of 94.4 %, while the 36 °C and control flasks were 97.9% and 97.4% viable respectively. This goes on to show that the control set point was best suited for the process and the 38 °C was the least favorable.

After this, a Student’s t-test was performed for each pair of set points to test for variance within the set points. The significance level *α* was set to 0.05. **Supplementary Figure 1** shows a plot of the means of the data from all set points following the t-test. The VCD didn’t seem to show significant difference across the set points in the t-test results though 38 °C showed the lowest mean VCD. Mean %Viability was severely affected in flasks incubated at a set point of 38 °C. It was seen that the control and 36 °C cultures showed significant similarities. The culture grown at 38 °C was significantly different than the other two set points.

Then, a multivariate analysis of all the responses in the data was performed. **Figure 3** shows a scatterplot matrix summarizes the bivariate relationships between each pair of response variables using the correlation coefficient *r*. The values of r in the correlation matrix range from (+1 = full correlation), (0 = no correlation), to (−1 = full inverse correlation) which can also be observed from the shape of the ellipse. We can see that there is no correlation whatsoever between the VCD and %Viability. However, VCD shows a strong inverse correlation with osmolality, glucose, glutamate, and pO_2_ and a positive correlation with lactate and pCO_2_ indicating that these responses highly covary and can be considered for dimensionality reduction in any further analysis. %Viability shows no correlation with any of the responses.

**Figure 3.**
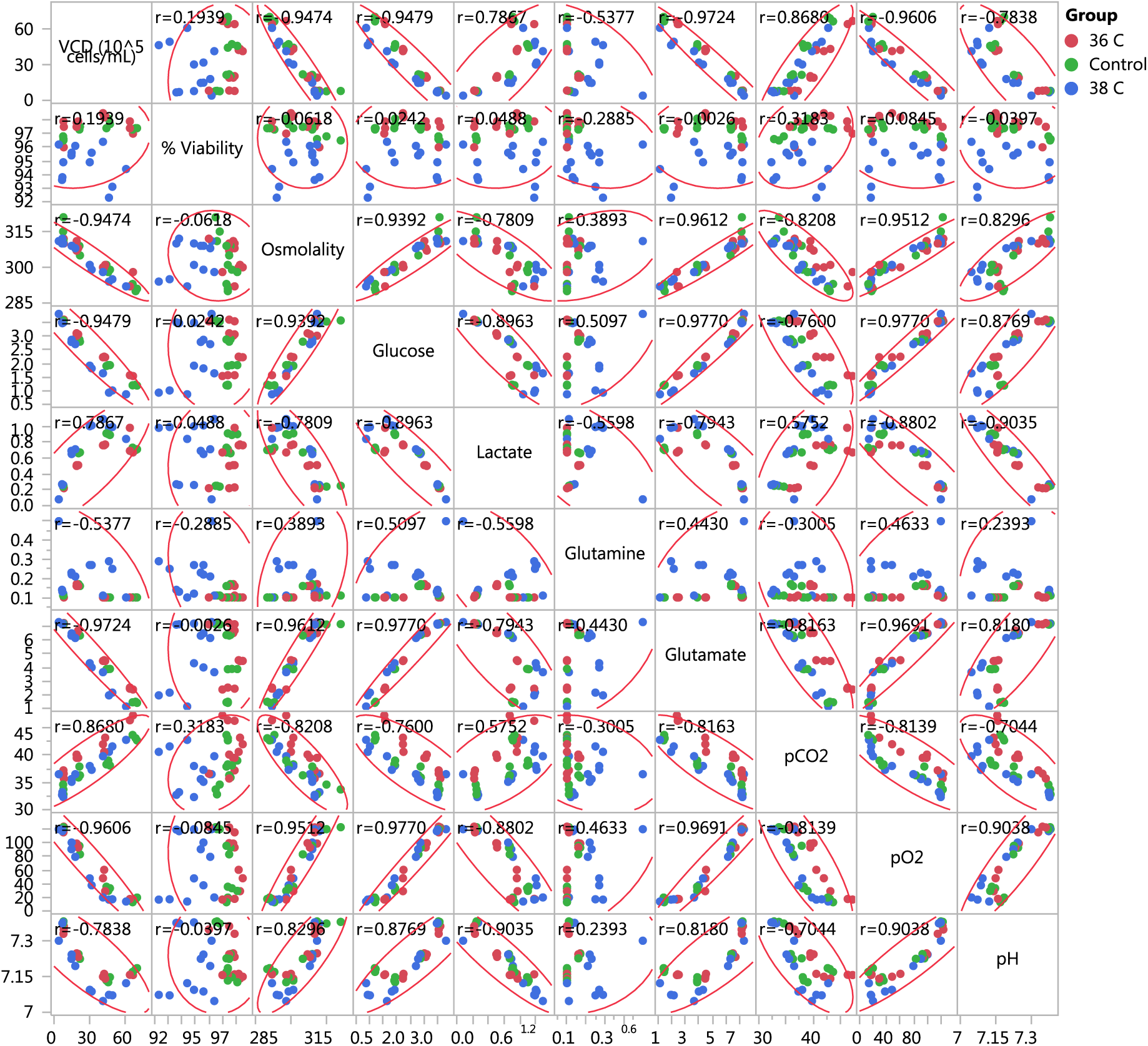
Multivariate analysis of all responses affecting culture growth. The ellipses in the scatterplot matrix depict the strength of the linear relationship between each pair of responses (variables) based on the correlation coefficient *r*. Each ellipse assumes that each pair of variables follows a bivariate normal distribution, and encloses approximately 95% of the points. The narrowness of the ellipse reflects the degree of correlation of the variables: If the ellipse is narrow and diagonally oriented, the variables are correlated and if the ellipse is fairly rounded and not diagonally oriented, the variables are uncorrelated. Also, the closer the value of r to +/-1 the more correlated the responses are. ^13^ VCD shows a strong inverse correlation with osmolality, glucose, glutamate, and pO_2_ and a positive correlation with lactate and pCO_2_. %Viability shows no correlation with any of the responses.

To assess the growth of the cells in the culture we plotted the glucose consumption and the lactate produced in all the flasks during the course of the experiment. Overall, the faster the glucose is consumed the more lactate is produced; however, this does not correlate to faster growth. Specifically, the flask incubated at 38 °C utilized glucose at the fastest rate and correspondingly, they also grew the poorest. As shown in **Figure 4**, the flasks incubated at higher temperatures are less efficient at converting glucose to energy. Though the control flasks were slower at utilizing the glucose than the flasks incubated at 36 °C, they showed higher VCD levels.

**Figure 4.**
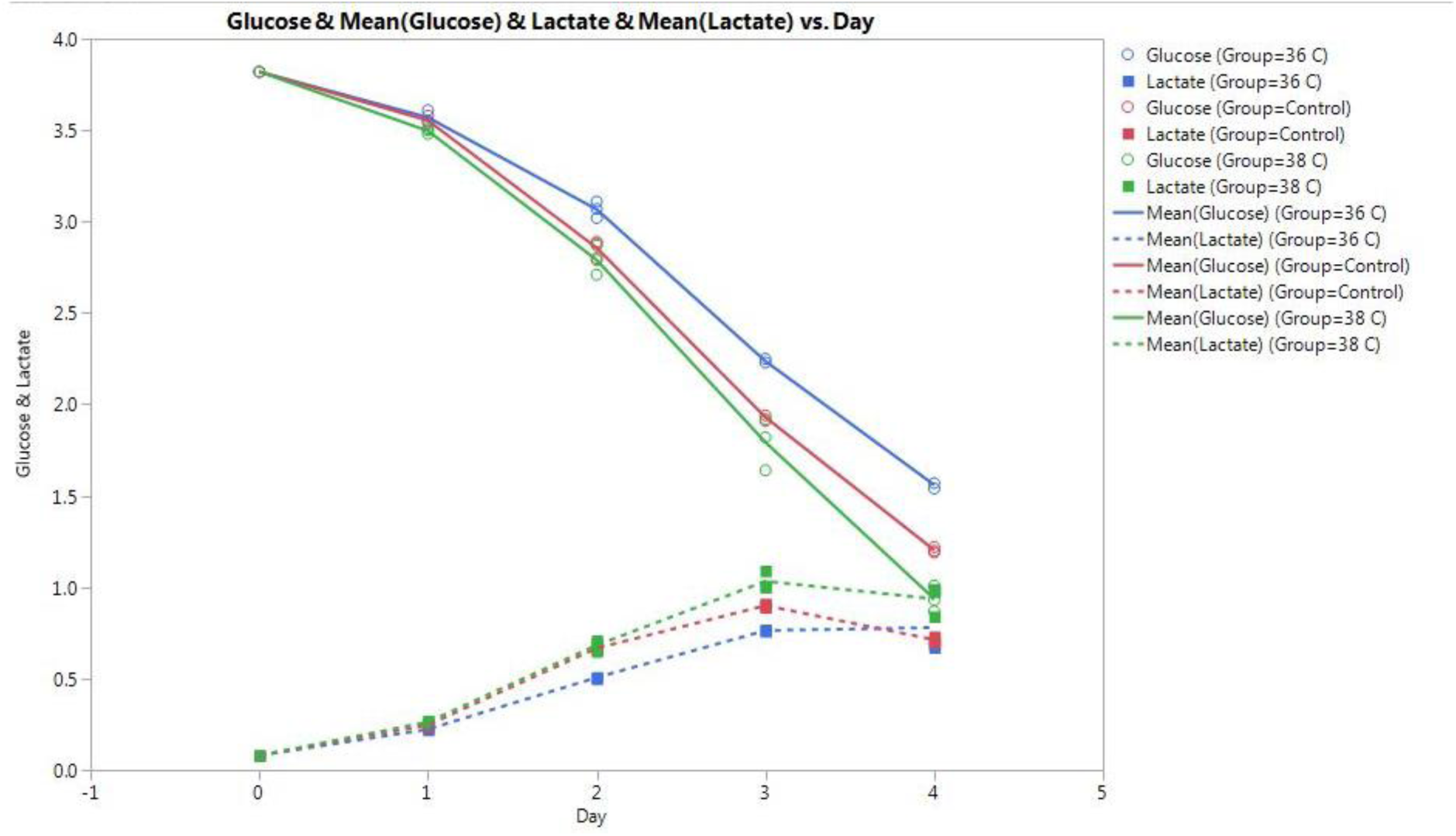
Effect of temperature on Glucose and Lactate levels in culture. The plot shows how the glucose and lactate levels are affected by the different temperature set points. We can see that the flasks incubated at 38 °C show the fastest consumption of glucose and had the poorest growth. The control flasks consumed glucose at slower rates and had slower lactate accumulation than the flask at 38 °C even though they reached the highest VCDs among all 3 set points. Even though the 36 °C flask had the slowest glucose consumption and lactate formation it had lower VCD than the control flask. This shows that high temperatures are most detrimental to the growth and the control conditions are most optimal for cell growth.

This shows that the control flasks consumed glucose at slower rate and had slow lactate accumulation, even though they reached the highest VCDs and %Viability during the course of the experiment.

### Effect of agitation rate on cell growth

As compared to the temperature set points the variations in agitation rate seemed to have lower impact on the VCD and viability of the culture. As shown in **Figure 5**, a least squares model was fitted for the data from all the flasks. The R^2^ value of 1 and 0.92 for VCD (**Fig. 5a**) and %Viability (**Fig. 5b**) respectively suggests that the model is a good fit for the data. It is also seen that all the points lie well along the 45 degree diagonal line which again indicates goodness-of-fit.

**Figure 5.**
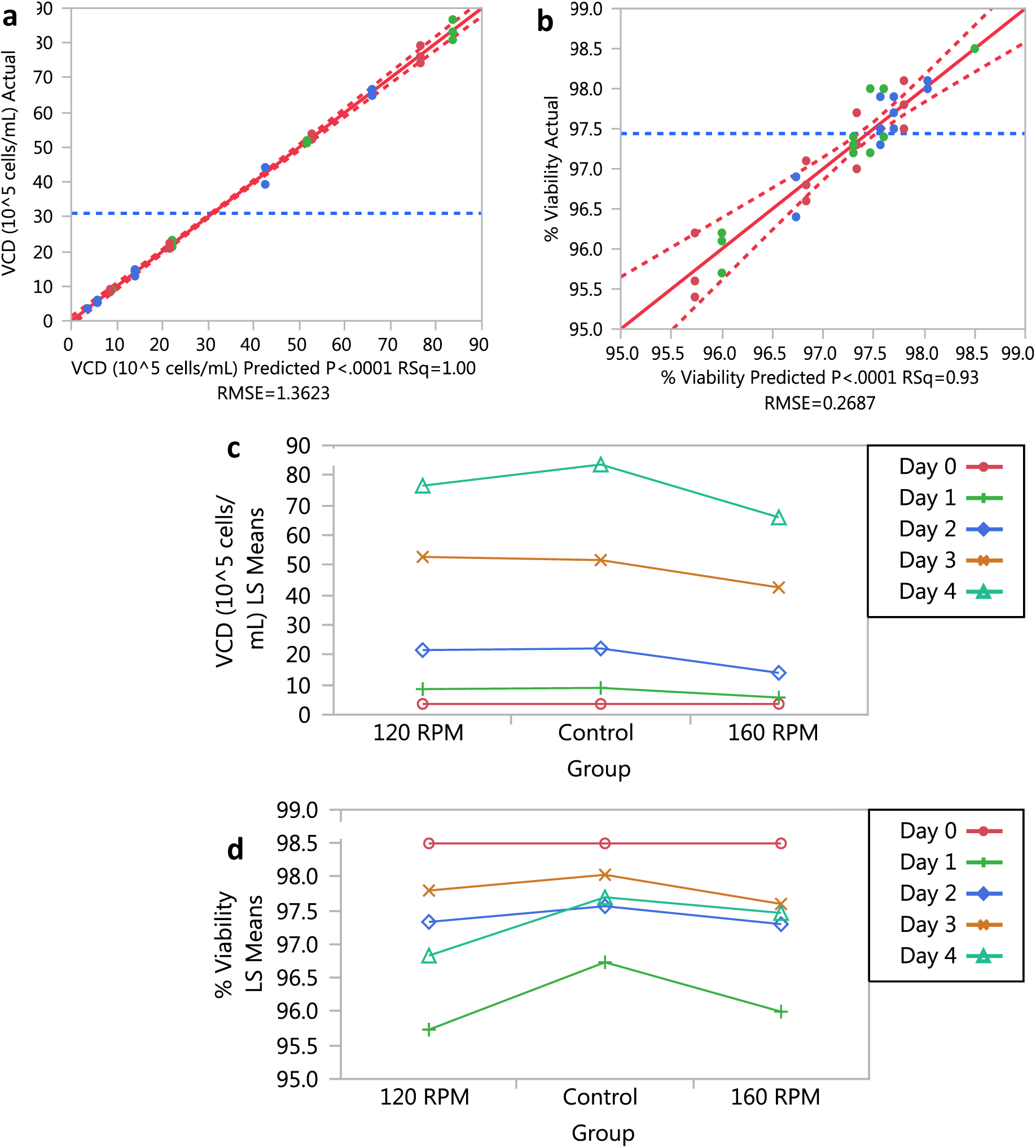
After fitting the linear model using regression analysis in JMP we need to determine if the model fits the data well. The results observed from the analysis can then be used to view any interactions between the responses. **(a,b)** Actual by Predicted plot. The observed values are plotted on the y-axis and the jackknife predicted values of the responses (effects) are on the x-axis for VCD as shown in **a** and %Viability as shown in **b**. This is the leverage plot for the whole model. We can predict the goodness-of-fit for the model by seeing how well all the points lie along the 45 degree line. The closer the value of R^2^ to 1 the better the model fits the data. As both these conditions were satisfied both the models were adjudged to be a good fit for the data. **(c,d)** Least Squares Means (LSMeans) plot. The least squares means for nominal and ordinal main effects and interactions were plotted after fitting the model in JMP. The results showed that in **c**, the VCD levels were not much affected by variation in the agitation rate. Over the duration of 4 days, the control group seemed to show the highest viable cell density indicating that the control agitation rate was best suited for the cell growth. At the end of Day 4 lowest VCD was observed in the flask set at 160 RPM. Similar results were observed in **d**, and at the end of Day 4 control showed the highest %viability while the 120 RPM showed the lowest %Viability. At the end of Day 4 however, both the 160 RPM and control flasks showed nearly the same values for %Viability. The 120 RPM set point proved to be the most unfavorable for cell growth.

Again, we calculated the Least Squares values from the model to view the results of the analysis. It showed the effects of all three agitation rate set points on the viable cell density across the span of 0-4 Days. It was observed that the control group seemed to show the highest viable cell density indicating that the control agitation rate was best suited for the cell growth. At the end of Day 4 lowest VCD was observed in the flask set at 160 RPM (**Fig. 5c**). Similar results were observed in the Least Squares Means (LSMeans) plot for %Viability; at the end of Day 4 control flasks seemed to show the highest %Viability as compared to the other flasks.

Both the 160 RPM and control flasks showed nearly the same values for %Viability. The 120 RPM set point was demonstrated to be the most unfavorable for cell viability.

The flask incubated at 160 RPM reached a maximum VCD of 66.66 × 10^5^ cells/mL as compared to a VCD of 79.27 × 10^5^ cells/mL in the 120 RPM flask and 86.73 × 10^5^ cells/mL in the control flask. After 4 days, the 120 RPM flask had a maximum viability of 96.8%, while the 160 RPM and control flasks were 97.2% and 97.9% viable respectively. This goes on to show that the control set point was best suited for the process and the other two set points were not as favorable. After this, a Student’s t-test was performed for each pair of set points to test for variance within the set points. The significance level *α* was set to 0.05.

**Supplementary Figure 2** shows a plot of the means of the data from all set points following the t-test. Both the VCD and %Viability didn’t show a significant difference across the set points in the t-test results although control flasks consistently showed higher values across both VCD and %Viability.

Then, a multivariate analysis of all the responses in the data was performed. **Figure 6** shows a scatterplot matrix summarizes the bivariate relationships between each pair of response variables using the correlation coefficient *r*. The values of r in the correlation matrix range from (+1 = full correlation), (0 = no correlation), to (−1 = full inverse correlation) which can also be observed from the shape of the ellipse. We can see that there is no correlation whatsoever between the VCD and %Viability. However, VCD shows a strong inverse correlation with osmolality, glucose, glutamate, and pO_2_ and a positive correlation with lactate and pCO_2_ indicating that these responses highly covary and can be considered for dimensionality reduction in any further analysis. %Viability shows no correlation with any of the responses.

**Figure 6.**
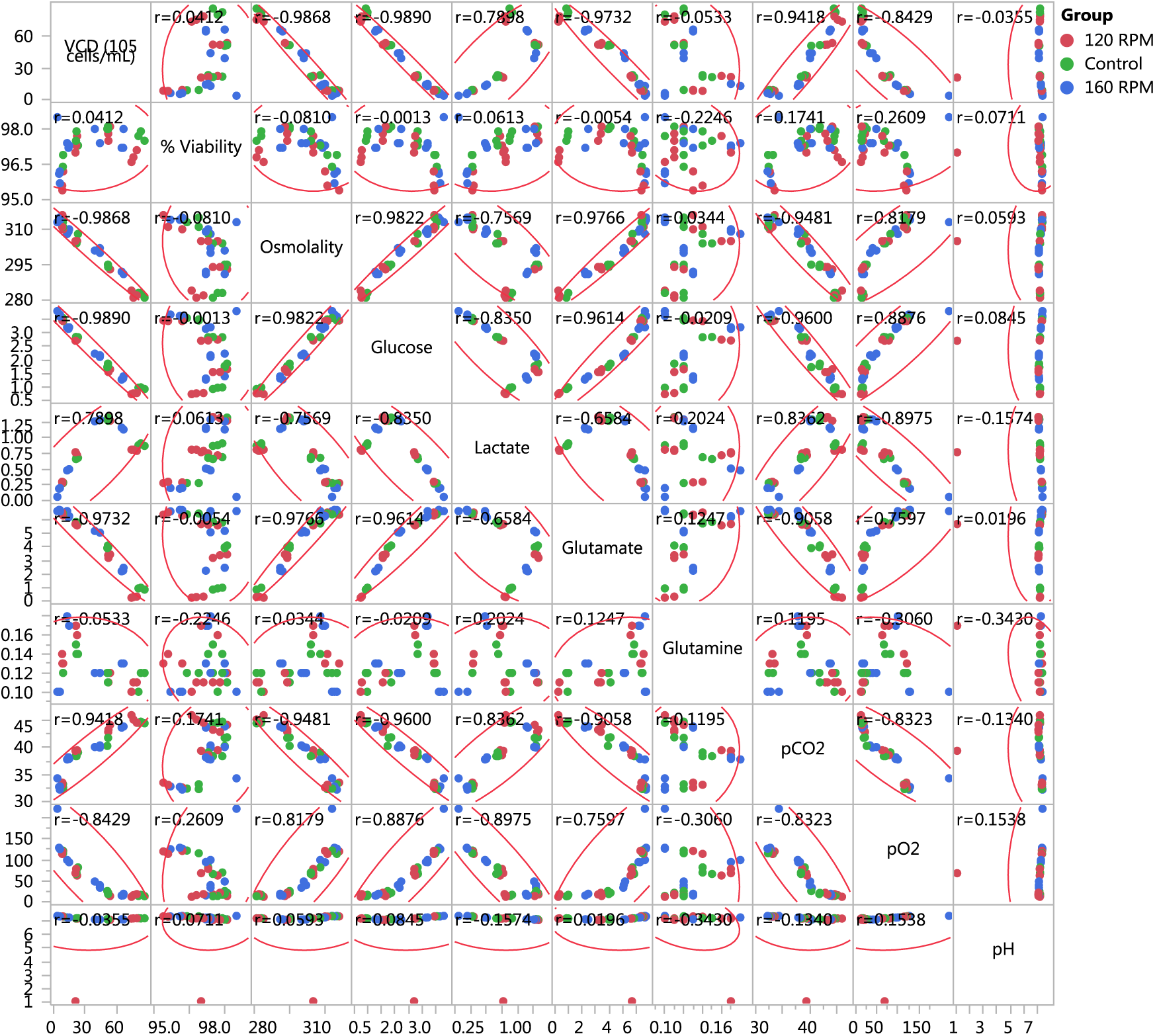
Multivariate analysis of all responses affecting culture growth. The ellipses in the scatterplot matrix depict the strength of the linear relationship between each pair of responses (variables) based on the correlation coefficient *r*. Each ellipse assumes that each pair of variables follows a bivariate normal distribution, and encloses approximately 95% of the points. The narrowness of the ellipse reflects the degree of correlation of the variables: If the ellipse is narrow and diagonally oriented, the variables are correlated and if the ellipse is fairly rounded and not diagonally oriented, the variables are uncorrelated. Also, the closer the value of r to +/-1 the more correlated the responses are. ^13^ VCD shows a strong inverse correlation with osmolality, glucose, glutamate, and pO_2_ and a positive correlation with lactate and pCO_2_. %Viability shows no correlation with any of the responses.

To assess the growth of the cells in the culture we plotted the glucose consumption and the lactate production in all the flasks during the course of the experiment. Overall, the faster the glucose is consumed the more lactate is produced; however, this does not correlate to faster growth. The flask incubated at 120 RPM utilized glucose at the fastest rate however, showed poorer growth than control flasks. As shown in **Figure 7**, though the control flasks were slower at utilizing the glucose than the flasks incubated at 120 RPM, they reached the highest VCDs among all 3 set points. This shows that the control flasks set point of 140 RPM was most ambient for culture growth.

**Figure 7.**
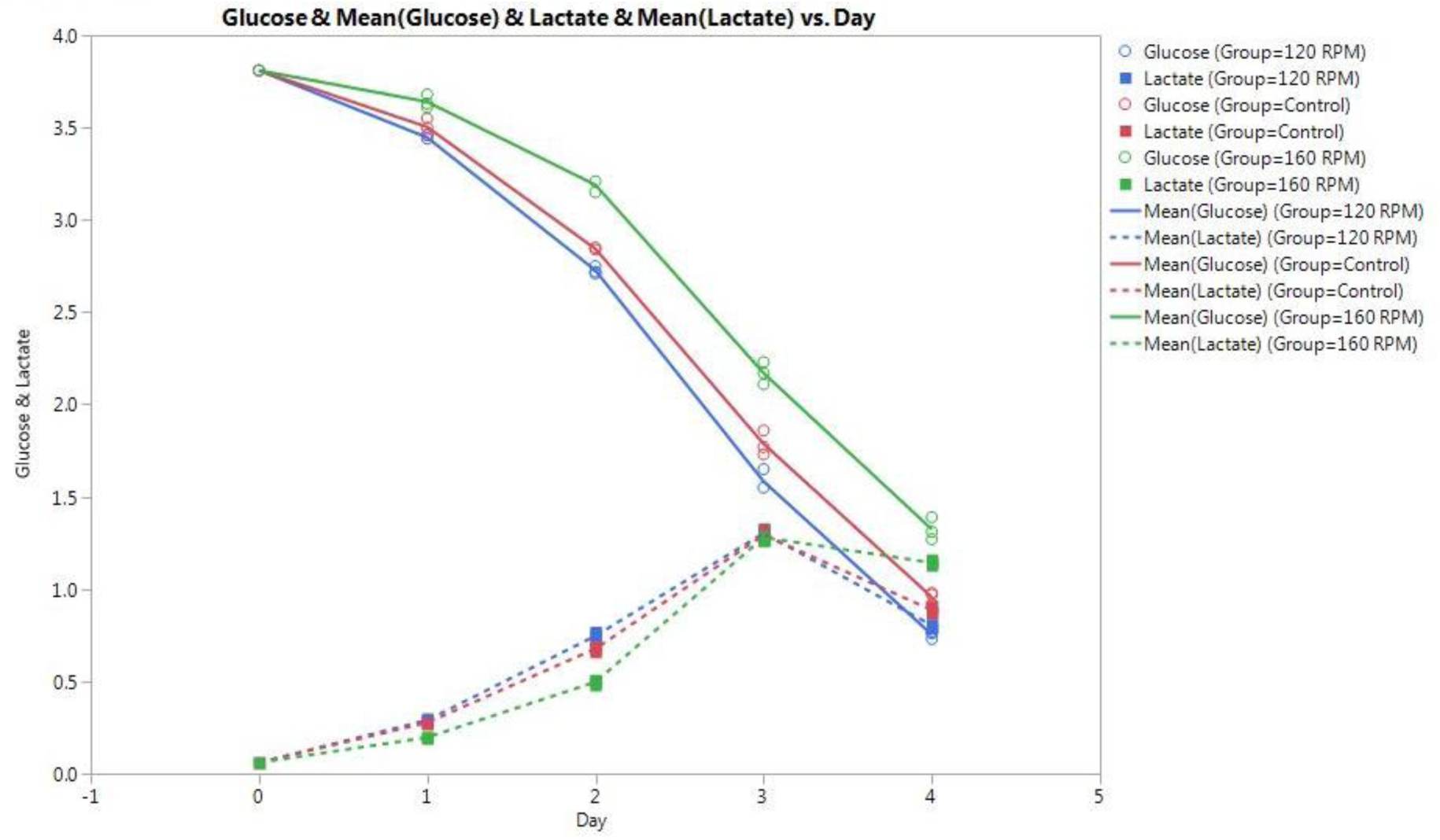
Effect of agitation rate on Glucose and Lactate levels in culture. The plot shows how the glucose and lactate levels are affected by the different RPM set points. We can see that the flasks incubated at 120 RPM show the fastest consumption of glucose but showed poorer growth than control flasks. The control flasks consumed glucose at slower rates and had slower lactate accumulation than the flask at 120 RPM even though they reached the highest VCDs among all 3 set points. Even though the 160 RPM flask had the slowest glucose consumption it had lower VCD than the control flask. This shows that the control conditions are most optimal for cell growth.

## DISCUSSION

A summary of the data from both the treatments is given in **Table 1**. We can see that overall control flasks show the most consistent results.

**Table 1.**
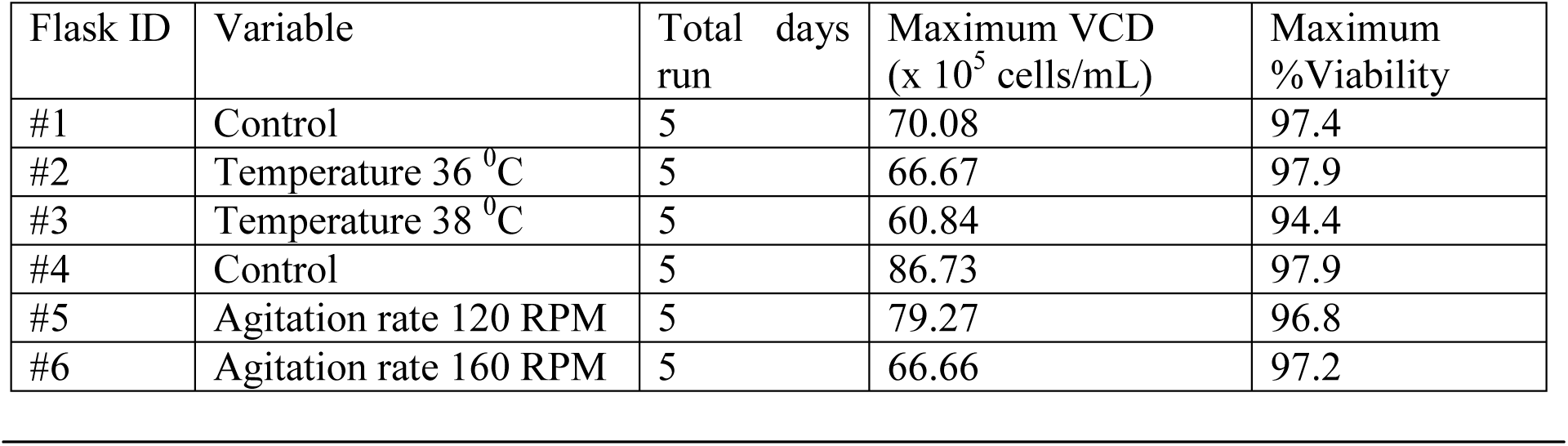
Summary of final data.

The individual effects of temperature and agitation rate on specific cell growth rate were evaluated in flask stage. Temperature has a greater effect on VCD and %Viability than agitation rate. Incubation at 38 °C is detrimental to NS0 cell culture growth. Incubation at varying agitation rates of 120 RPM and 160 RPM had minimal effect on the growth of cultures. Incubation at 37 °C and 140 RPM appears to be the optimal set points for growth in shake flask conditions.

These results show that through a suitable experimental approach it is possible to evaluate the differential effect of temperature and agitation rate modulations on the production of monoclonal antibodies in NS0 cells, showing that the cellular response in the studied system is primarily modulated by changes in the culture temperature and to a lesser extent by changes in the agitation rate.

Also, extrapolating results from the small-scale stage to the production stage might not always yield accurate predictions of the process data. Hence, further research for assessing the implications of critical process parameters on advanced cell culture stages is required. Understanding how these variables individually affect the different stages of bioprocessing is a critical aspect on which future efforts should be focused.

## MATERIALS AND METHODS

### Cell line

A GS-NS0 murine myeloma cell line expressing an IgG1λ monoclonal antibody (mAb) was used in this research. The NS0 Clone#97 was obtained from the GSK WCB-P.5.

### Media preparation

The seed culture medium used was CD Hybridoma AGT medium, supplemented with HGS dried powder medium, glucose and amino acids. The seed medium was also supplemented externally with cholesterol as the NS0 cell lines used are cholesterol-dependent auxotrophs. ^10^

### Flask inoculation and incubation

Cell culture experiments were performed in 250 ml shake flasks maintaining a working volume of 100 ml with a seeding density of 4 × 10^5^ cells/mL. The flasks were inoculated and maintained in the incubator for 4 days.

A series of two experiments was performed, in triplicate, at 36 °C and 38 °C and at 120 RPM and 160 RPM. The controls were maintained as triplicates at 37 °C and 140 RPM for both the experiments. **Table 2** shows the design followed for the experimental setup.

**Table 2.**
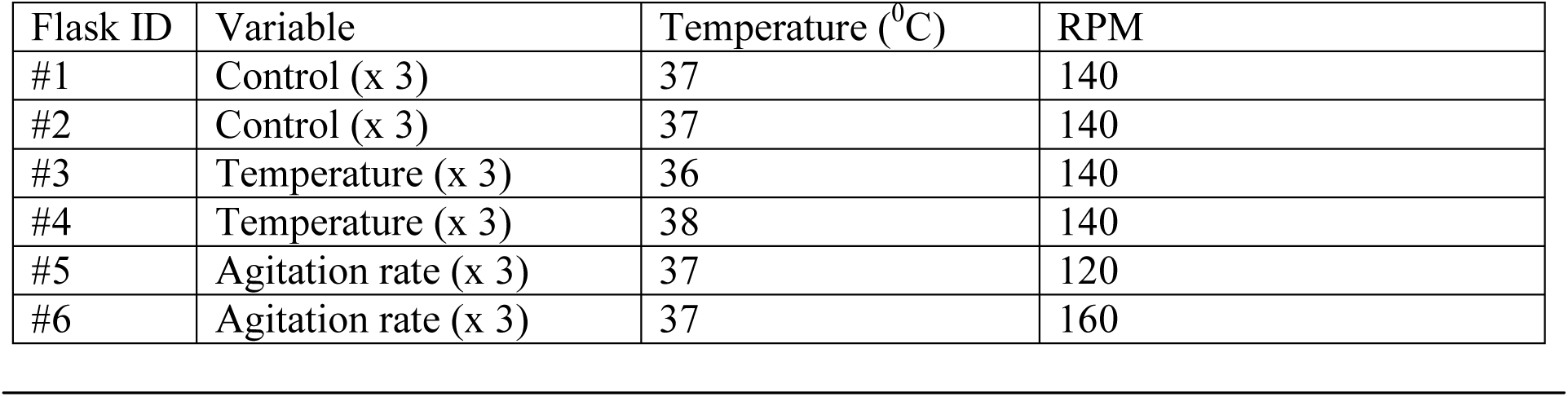
Incubation conditions and flask details.

### Flask sampling

Shortly after inoculation and every 24 h thereafter experimental flasks were sampled. Samples were taken every 24 +/-2 hours for viable cell quantification and analytical measurement separately. Each day samples were analyzed for VCD, %viability, pH, pCO_2_, pO_2_, glucose, lactate, glutamine and glutamate.

### Analytical methods

**Table 3** shows a summary of the assays performed for the analysis of the samples.

**Table 3.** Daily Bioreactor Sample In-Process Assays.

| Instrument | Assay |
| --- | --- |
| ViCell | Viable Cell Density ($\times 10^5$ cells/mL) |
|  | Viability (%) |
| | Average Cell Diameter ($\mu\text{m}$) |
| Blood Gas Analyzer (BGA) | Offline pH |
|  | Offline pO<sub>2</sub> (mmHg) |
|  | Offline pCO<sub>2</sub> (mmHg) |
| Cedex Bio HT | D-Glucose (g/L) |
|  | L-Lactate (g/L) |
|  | L-Glutamine (mmol) |
|  | L-Glutamate (mmol) |
| Micro Osmometer | Osmolality (mmol/ kg) |

Cell viability and Viable Cell Density (VCD) were determined using a Vi-CELL Cell Viability Analyzer (Vi-CELL XR cell counter, Beckman Coulter).

The pO_2_, pCO_2_ and pH were measured using a Blood Gas Analyzer (248 BGA, Chiron Diagnostics).

Osmolality was measured using a Micro Osmometer (2020 Multi-Sample Micro Osmometer, Advanced Instruments Inc.).

Residual glucose, lactate, glutamine and glutamate concentrations were determined with an automatic Select Biochemistry Analyzer (Cedex Bio HT, Roche Inc.). All samples were micro-centrifuged for 1 minute at 16000 RPM prior to analysis and the undiluted supernatant was immediately used for analytical measurements.

### Statistical analysis

The cell cultures at each condition were performed in triplicates and one independent sample was taken at each time point for every culture from each flask with analytical measurements carried out separately.

Bivariate and Multivariate analysis of all the factors and responses and analysis of variance was used to compare the results using JMP^®^ 12 software (SAS Institute Inc.) for Windows. ^11^

## ACKNOWLEDGMENTS

The work was entirely conducted in the GlaxoSmithKline Biopharmaceutical Technology Lab under the supervision of Yuan Tian. The opportunity to conduct the research and the guidance provided by Yuan Tian is appreciated and the support received from Samruddhi Patil for sample analysis is recognized. Buffy Hudson-Curtis assisted significantly with the statistical data analysis.

## SUPPLEMENTARY INFORMATION

This section highlights the supplementary figures from the paper. The scripts that were developed to perform the data analysis of the experimental results for the research project were primarily written in JMP Scripting Language or JSL. Special emphasis was given to data reproducibility for all the figures listed in the paper. Original scripts and source data can be obtained from the author at any time upon request.

**Supplementary Figure 1.**
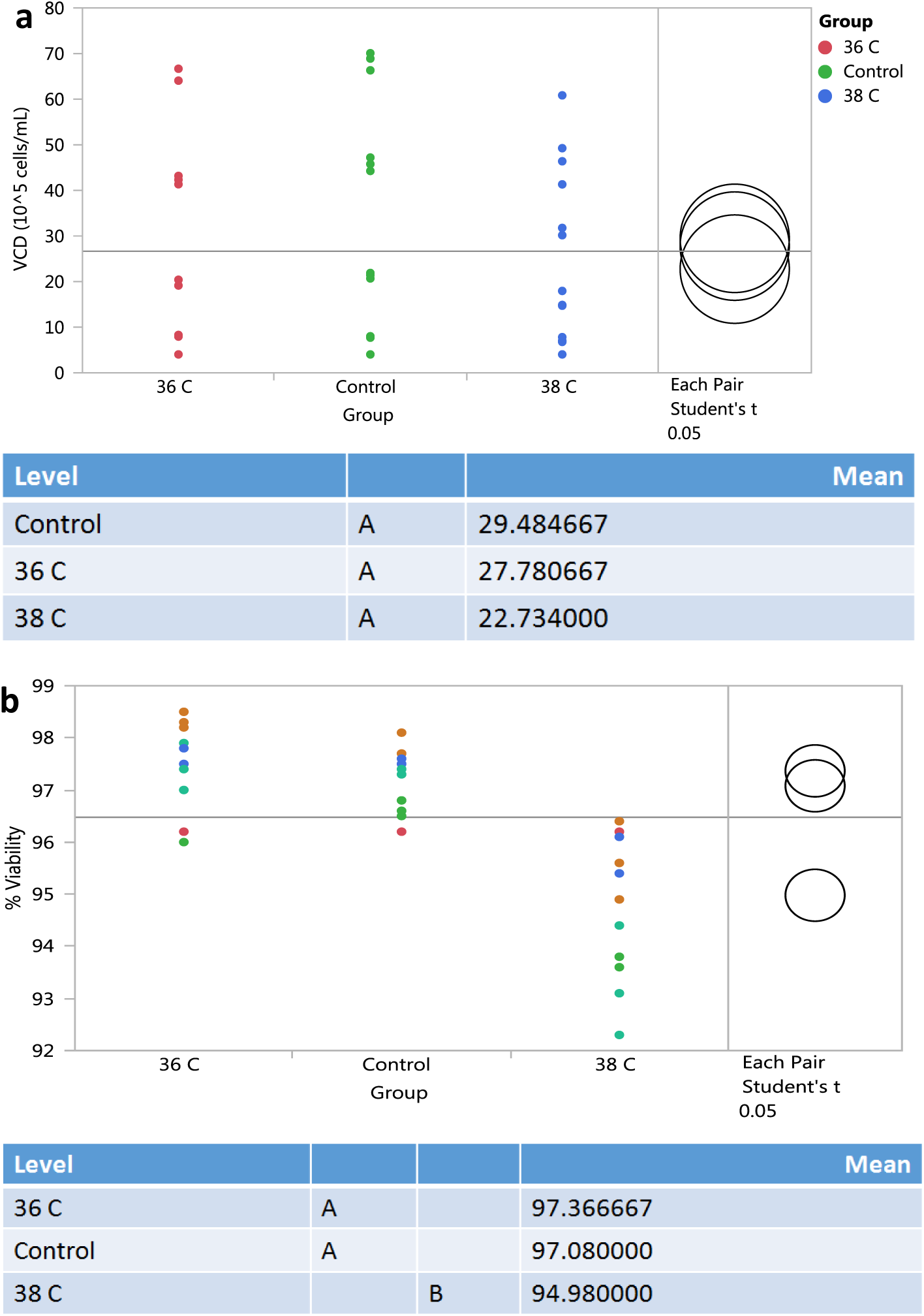
A plot of the means of the data from all temperature set points following the t-test. The levels not connected by the same letter are significantly different. **(a)** The VCD didn’t seem to show significant difference across the set points in the t-test results though 38 °C showed the lowest mean VCD. **(b)** Mean %Viability was also severely affected in flasks incubated at a set point of 38 °C. It was seen that the control and 36 °C cultures showed significant similarities than the culture grown at 38 °C.

**Supplementary FIgure 2.**
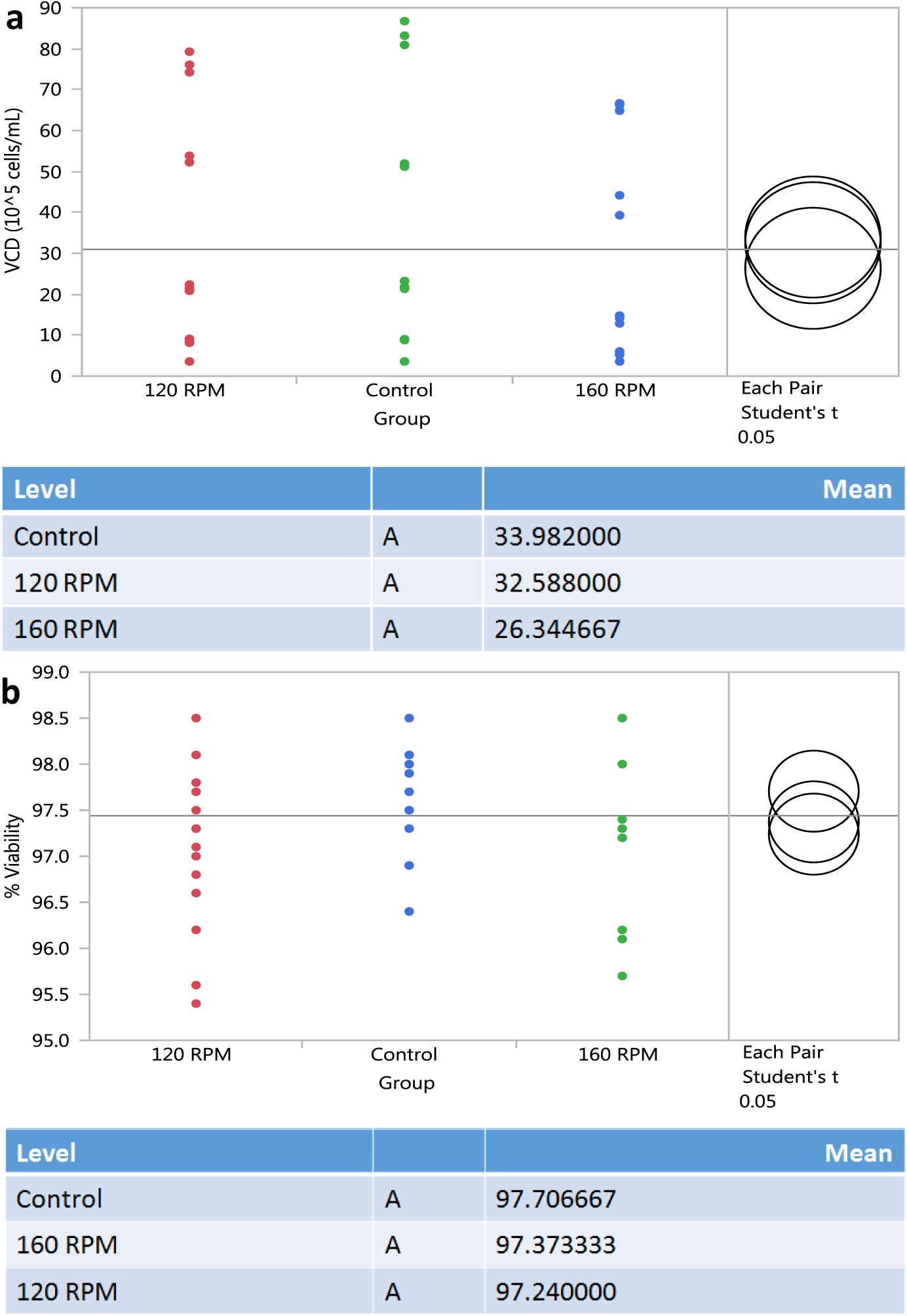
A plot of the means of the data from all set points for agitation rate following the t-test. The levels not connected by the same letter are significantly different. **(a)** The VCD didn’t seem to show significant difference across the set points in the t-test results though 160 RPM showed the lowest mean VCD. **(b)** Mean %Viability was also seen to show no significant difference between the set points. It was seen that the control flasks showed consistently higher mean values.

